# Identifying transcription patterns of histology and radiomics features in NSCLC with neural networks

**DOI:** 10.1101/2020.07.22.215558

**Authors:** Nova F. Smedley, Denise R. Aberle, William Hsu

## Abstract

**Purpose:** To investigate the use of deep neural networks to learn associations between gene expression and radiomics or histology in non-small cell lung cancer (NSCLC).

**Materials and Methods:** Deep feedforward neural networks were used for radio-genomic mapping, where 21,766 gene expressions were inputs to individually predict histology and 101 CT radiomic features. Models were compared against logistic regression, support vector machines, random forests, and gradient boosted trees on 262 training and 89 testing patients. Neural networks were interpreted using gene masking to derive the learned associations between subsets of gene expressions to a radiomic feature or histology type.

**Results:** Neural networks outperformed other classifiers except in five radiomic features, where training differences were <0.026 AUC. In testing, neural networks classified histology with AUCs of 0.86 (adenocarcinoma), 0.91 (squamous), and 0.71 (other); and 14 radiomics features with >= 0.70 AUC. Gene masking of the models showed new and previously reported histology-gene or radiogenomic associations. For example, hypoxia genes could predict histology with >0.90 test AUC and published gene signatures for histology prediction were also predictive in our models (>0.80 test AUC). Gene sets related to the immune or cardiac systems and cell development processes were predictive (>0.70 test AUC) of several different radiomic features while AKT signaling, TNF, and Rho gene sets were each predictive of tumor textures.

**Conclusion:** We demonstrate the ability of neural networks to map gene expressions to radiomic features and histology in NSCLC and interpret the models to identify predictive genes associated with each feature or type.

**Author Summary:** Non-small-cell lung cancer (NSCLC) patients can have different presentations as seen in the CT scans, tumor gene expressions, or histology types. To improve the understanding of these complementary data types, this study attempts to map tumor gene expressions associated with a patient’s CT radiomic features or a histology type. We explore a deep neural network approach to learn gene-radiomic associations (i.e., the subsets of co-expressed genes that are predictive of a value of an individual radiomic feature) and gene-histology associations in two separate public cohorts. Our modeling approach is capable of learning relevant information by showing the model can predict histology and that the learned relationships are consistent with prior works. The study provides evidence for coherent patterns between gene expressions and radiomic features and suggests such integrated associations could improve patient stratification.

## 1 Introduction

Personalized medicine has driven the study of high-throughput profiling of both molecular and medical imaging data to discern survival, treatment outcomes, and subtyping in cancer patients. In particular, radiogenomic studies attempt to integrate the two complementary data types to explain tumor imaging patterns using molecular information and vice versa. For example, radiogenomic studies have shown molecular patterns (e.g., gene expression, gene mutation, or molecular subtype) can be predicted from imaging features (e.g., computed tomography (CT) or magnetic resonance (MR) images) [1, 2, 3, 4, 5, 6]. Radiogenomic studies support the derivation of tumors’ biological states from noninvasive imaging and the association mapping of molecular and imaging observations to better understand cancer heterogeneity. However, radiogenomic studies are often limited by the use of feature selection, linearity assumptions, and lack of testing datasets [7].

Deep learning techniques have been widely used on molecular and imaging datasets due to their ability to have high-dimensional input without the need for engineered features, and model nonlinear and hierarchical relationships. Several studies have used deep learning models such as convolutional neural networks, generative adversarial networks, and autoencoders for radiogenomic mapping [5, 8, 9, 6]. However, such works focus on accurate prediction of imaging features or imaging characteristics and often do not provide insight into the biological differences leading to accurate predictions by the models. While classification performance is a factor, it is also of interest to probe the trained models to validate the learned radiogenomic associations and to discover new hypotheses.

In our previous work, we addressed the model understanding challenge by presenting methods, such as gene masking to interpret trained neural networks [10]. We showed the models were capable of learning radiogenomic associations that were consistent with prior work while also generating new associations for further consideration. A limitation of [10] is that the analysis was performed using a single dataset from glioblastoma patients. We did not demonstrate the generalizability of our approach in other domains. Therefore, the purpose of this study was to investigate the ability of neural networks to model gene expression in different radiogenomic datasets, such as in a different cancer domain in which there are multiple different histologies or stage, there are computationally derived imaging features such as textures, and there is an external validation or testing dataset available.

Here, we present deep feedforward neural networks to model transcriptomes using two similarly derived radiogenomic datasets recently published in non-small cell lung cancer (NSCLC) [11]. As one of the few publicly released radiogenomic datasets available, the paper reported radiogenomic associations and provided a basis for comparison. First, we evaluate the ability of neural networks to separately predict each of two clinical traits and 101 radiomic features using a transcriptome consisting of 21,766 gene expressions in a training dataset of 262 patients. Next, we demonstrate the generalizability of our neural network models in the external dataset of 89 patients. Finally, we systematically probe the trained neural networks to define specific patterns of gene expression related to a clinical trait or radiomic feature. We compare the model’s learned relationships to previously published associations in similar work.

## 2 Materials and methods

### 2.1 Data

All transcriptomes, radiomics, and clinical data were downloaded from [11], see Supp. Table S1. `Dataset1` (262 patients from a North American institution) and `Dataset2` (89 patients from a European institution) were used as the training and testing datasets. Briefly, each transcriptome was measured on the same Affymetrix microarray chip and listed 21,766 gene expressions. Clinical stage referred to pathologic TNM staging and was defined as I, II, III or other. Pathologic histology was defined as adenocarcinoma, squamous carcinoma, or other. Radiomic features were extracted from three-dimensional tumor volumes in contrast-enhanced presurgical CT scans to determine histogram statistics; morphology; textures, such as gray-level co-occurrence matrix (GLCM), gray-level run-length matrix (GLRL, called RLGL), gray-level size-zone matrix (GLSZM); Laplacian of Gaussian (LoG) transformations; and wavelet decompositions.

In this study, each radiomic feature was a continuous variable that was transformed into binary classes using k-means clustering in the training dataset. Dichotomizing patients into two groups allowed us to focus on broad differences in the cohort for this initial work. For example, surface area to volume ratio (one radiomic feature) was grouped into two clusters: cluster A had a mean ratio of 0.26, and cluster B had a mean ratio of 0.52. Each cluster represented one class. Clustering was performed via `KMeans` from `sklearn`. Radiomic features with class frequency of > 10% were kept and resulted in 101 radiomic features. Clusters in the testing were defined by the clusters created in training. Distributions and correlations for classes are shown in Supp. Figs. S1–S2. For more details about the radiomic features, see Supp. Methods and [11, 12, 13].

### 2.2 Radiogenomic modeling

Dense feed-forward neural networks were used to map transcriptomes (model inputs) to radiomic features (model outputs). Gene expression was standardized by mean subtraction and standard deviation division for each gene. Layers within the radiogenomic models were constructed with three or four hidden layers, where the first hidden layer had either 2000, 4000, or 6000 nodes. The number of hidden nodes was halved with each subsequent hidden layer. Neural networks were trained using cross-entropy loss, nonlinear activation functions, dropout, batch normalization, and early stopping by monitoring loss.

Performance was measured with the receiver operating characteristic curve via the area under the curve (AUC). Hyperparameters were grid searched and the hyperparameters with the highest mean AUC in 10-fold cross-validation in the training dataset were selected. In cross-validation, metrics for gene standardization were based on the training folds for each split; in testing, metrics were based on the entire training dataset. Training performances were averaged cross-validation folds. Accuracy was calculated at a threshold of 0.5 class probability.

Figure 1a depicts the overall procedure used to train radiogenomic neural networks. To reduce the hyperparameter search for each radiomic feature, the grid search was applied in a neural network that predicted a *group* of radiomic features simultaneously, based on type of radiomic feature. A group was defined as a set of related features; the seven different groups are shown in Supp. Figs. S2. For example, the model would take a patient’s transcriptome and predict all 22 GLCM features at once as a multi-label classification task. One neural network was trained for each radiomic group. The best performing hyperparameters for a group were then used in a neural network to predict a *single* radiomic feature within the group. Other classifiers, including logistic regression, support vector machines, random forest, and gradient boosted trees were also considered. Each alternate model was trained to predict a single radiomic feature and evaluated against a radiogenomic neural network. The best performing models for each radiomic feature classification were then retrained on the entire training dataset to obtain the final models. Final models were then evaluated with the testing dataset. Radiomic features with at least 0.70 test AUC were kept for further analysis.

**Figure 1:**
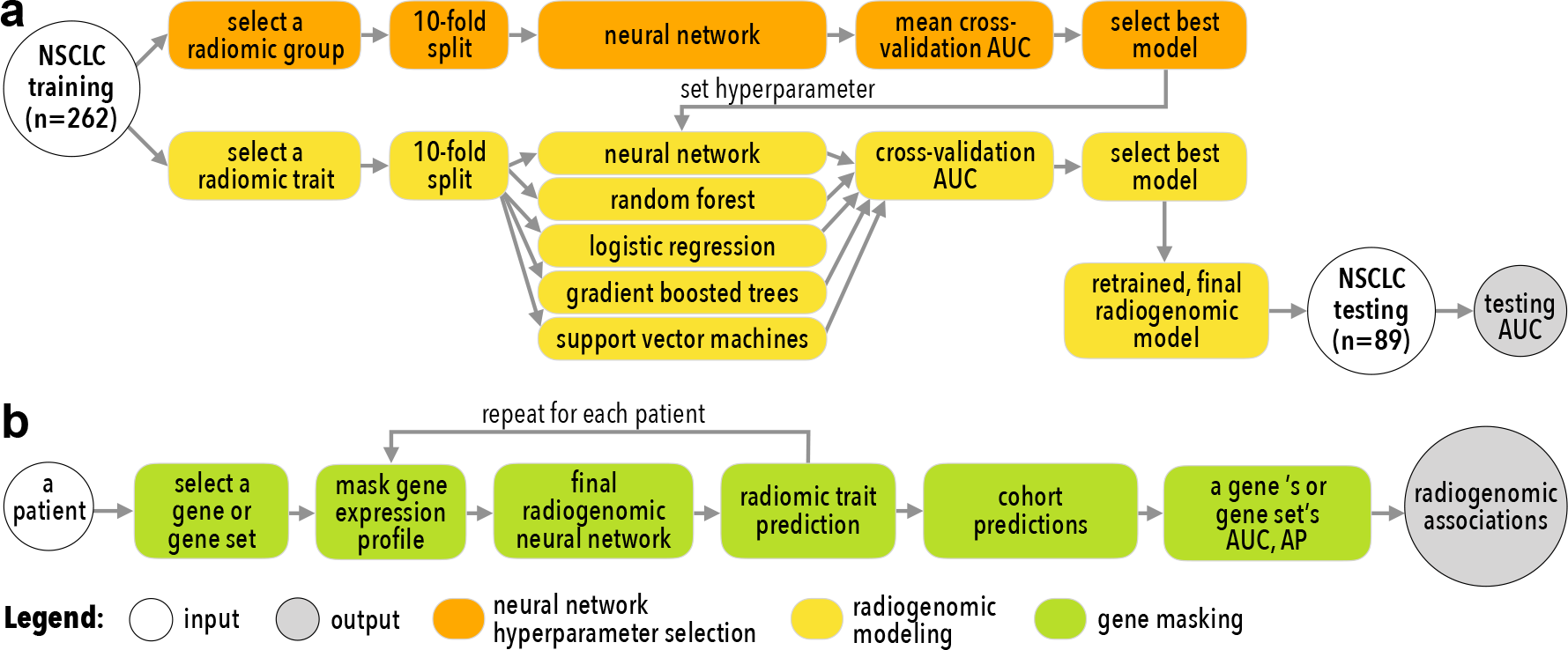
An overview of this study’s approaches to (**a**) training and (**b**) interpretation radio-genomic neural networks.

More details on the architecture of radiogenomic neural networks are shown in Supp. Fig. S3. All models and their hyperparameters are listed in Supp. Table S2.

### 2.3 Histology and stage modeling

Transcriptomes were also mapped to histology and stage in two additional neural networks constructed with a similar architecture as the radiogenomic neural networks. Instead of pre-dicting radiomic feature classes, models were trained to predict history or stage. Histology and stage were each modeled as multiclass classifications (Table S1), where the neural networks used categorical entropy loss and softmax activation in the prediction layer. Train scores were based on micro-averages across all classes and folds in cross-validation. Test scores and model interpretation methods were based on one class versus all other classes unless otherwise noted. The models were also trained using an approach similar to the radiogenomic modeling, namely the methods for hyperparameter and model selection via grid search in the training set and the comparison with other classifiers.

### 2.4 Extracting predictive gene expression patterns from neural networks

Figure 1b illustrates the gene masking steps used to extract predictive gene expression patterns from trained neural networks. Predefined gene sets were used for gene masking. These included the Hallmark and Gene Ontology (GO) biological processes from Molecular Signature Database (MSigDB) v7.0 [14]. Gene masking focused on a component of the model’s input by using one predefined gene set at a time, a process related to sensitivity analysis [15]. For example, masking of genes related to hypoxia involved taking each patient’s gene expression profile, keeping only the hypoxia genes’ expressions and setting all other gene expressions to zero (i.e., the input is masked). The masked input was then pushed through the model and the model’s prediction of a radiomic feature was recorded. This process was repeated for the entire cohort and classification performance was calculated. Gene masking measured the model’s ability to predict radiomic features based on gene expressions of a particular gene set, where the higher the performance the stronger the association in the cohort, as initially used in [10]. Masking resulted in “radiogenomic associations” learned by the model. The strength of the associations was by test AUCs, which represented how well a gene set could retain the classification performance in a radiogenomic neural network. For simplicity, gene sets with a maximum of 500 genes were studied.

Relationships between gene expression and histology were also studied using gene masking. Gene expressions were previously used to predict histology in NSCLC. In one work, a 42-gene signature was used to distinguished adenocarcinomas (ADC) from squamous cell carcinoma (SCC) [16]. In another similar effort, a 75-probe set signature was found for ADC, SCC, and large cell carcinoma (LCC) [17]. These two signatures were defined as four additional gene sets for comparison in this work.

## 3 Results

### 3.1 Predicting histology or stage from the transcriptome

#### 3.1.1 Neural networks outperform other models

In predicting histology and stage, most models achieved cross-validated AUC above 0.5; neural networks achieved the lowest error in the training dataset (Figure 2a). In testing, the neural network was unable to estimate stage with a test AUC equal to random guessing (0.5 AUC; Figure 2b). However, the histology model was able to generalize to the testing dataset and achieved the respective test scores of 0.86 AUC, 0.91 AUC, and 0.71 AUC in predicting adenocarcinomas, squamous cell carcinoma, or other histology, respectively; this resulted in an overall score of 0.77 test AUC when micro-averaged across classes. For this reason, the histology neural network was further analyzed.

**Figure 2:**
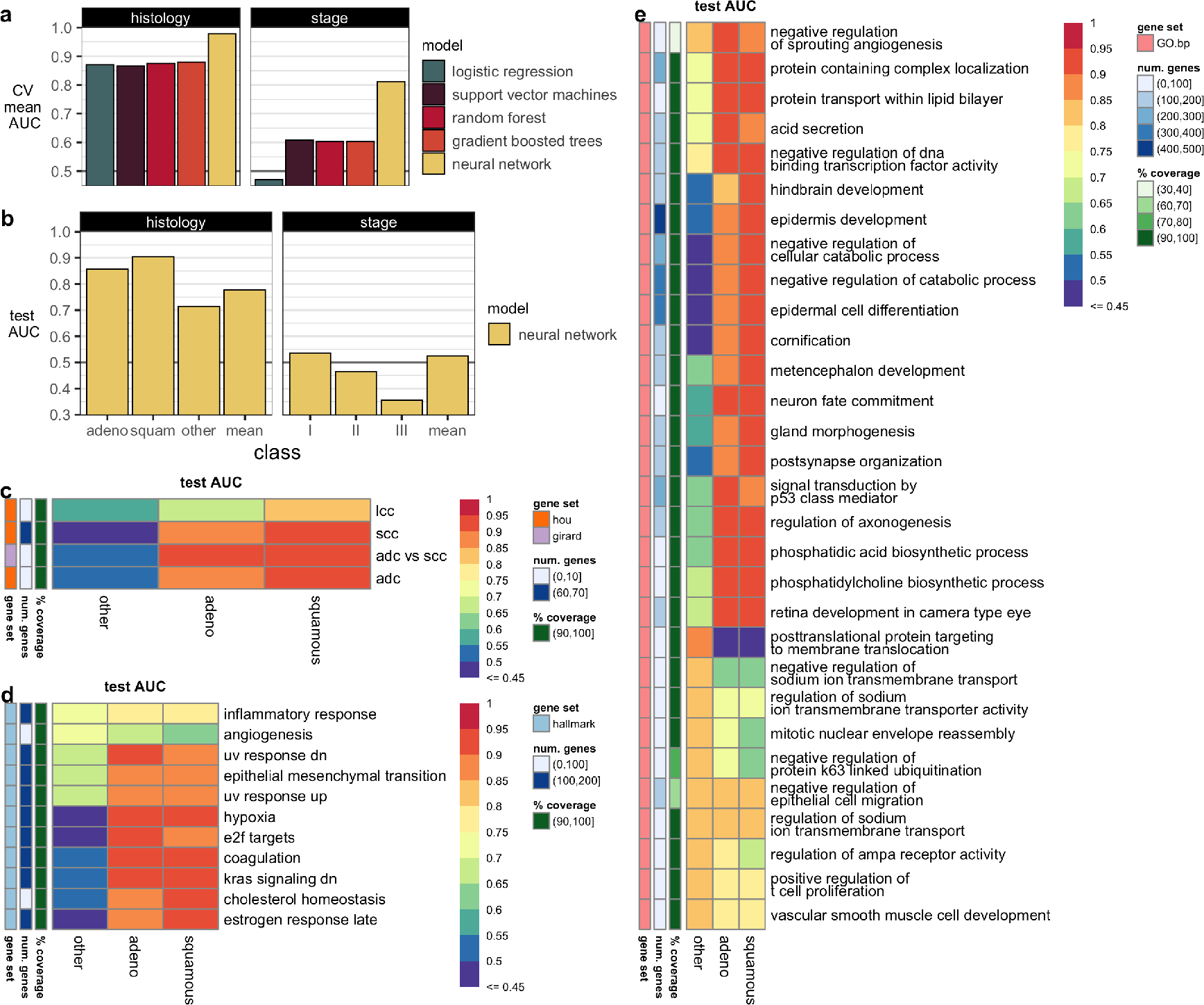
The ability of models to predict NSCLC histology and stage in (**a**) training and (**b**) testing. Gene masking of the histology neural network using gene sets from (**c**) published gene signatures for histology [16, 17], (**d**) Hallmarks (top five out of 50) [18], and (**e**) Gene Ontology biological processes (GO.bp, top ten out of 7350) [14]. CV=cross-validation.

#### 3.1.2 Gene masking identities gene sets that predict histology

Gene masking of the histology neural network showed agreement with previously published gene signatures [16, 17] for predicting NSCLC adenocarcinomas (ADC) and squamous cell carcinoma (SCC). As shown in (Figure 2c), the aforementioned gene signatures were also found to be predictive in our histology neural network in the testing dataset. In particular, the gene signature from [16] resulted in 0.93 test AUC in both ADC and SCC.

Hallmark gene sets were also indicative of histology (Figure 2d). Gene expressions related to hypoxia, coagulation, and KRAS signaling could predict both ADC and SCC (>0.90 test AUC). Similar to the overall test performances observed in Figure 2b, the histology neural network was driven by accurate predictions in ADC and SCC, where test performances were about 0.20 AUC higher than the “other histology” class. Subsequently, this behavior was reflected in gene masking, where the most predictive gene sets for estimating “other histology” in testing were inflammatory response (0.73 AUC) and angiogenesis (0.72 AUC). Notably, angiogenesis was more predictive of other histologies (0.73 AUC) than ADC (0.66 AUC) or SCC (0.63 AUC) classes in testing.

The most predictive GO biological processes (Figure 2e) were also associated with angiogenesis, epithelial mesenchymal transition, and hypoxia from the Hallmark gene sets. Of the gene sets considered in gene masking, `negative regulation` of `DNA-binding transcription factor activity` from GO had the best overall testing performance with the highest micro-averaged AUC of 0.79; the individual test AUCs were 0.93 in ADC, 0.91 in SCC, and 0.75 in other. The aforementioned gene set consisted of 170 genes, where 156 (91.2%) were in the gene expression profile. A summary of gene expression patterns is in Table 1. Notably, *KRAS* is a major gene studied in NSCLC and the KRAS Hallmark was found to be predictive of ACC and SCC.

**Table 1:**
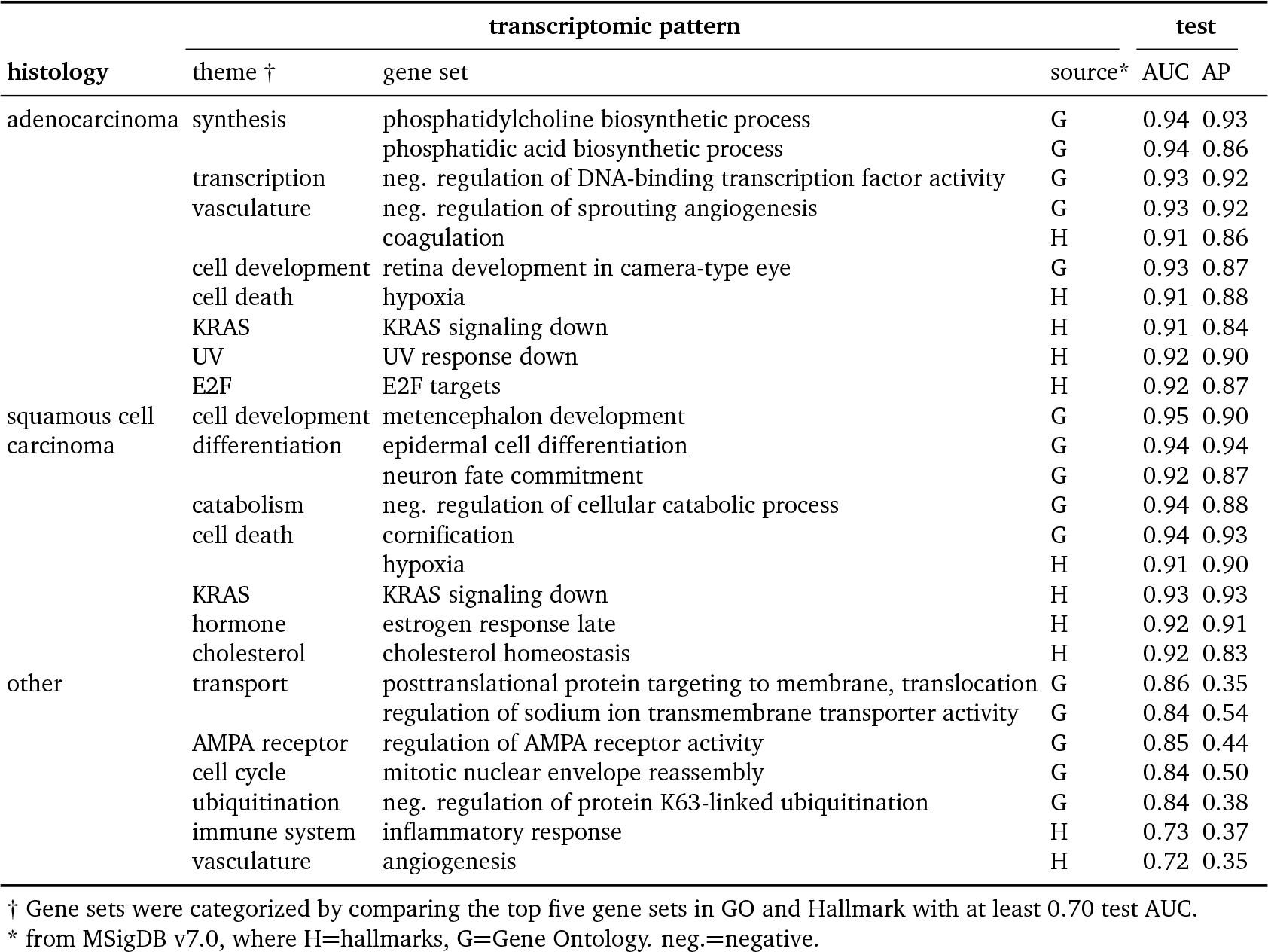
Summary of predictive gene expression patterns in NSCLC histology.

### 3.2 Predicting CT imaging features from the transcriptome

#### 3.2.1 Performance

Neural networks were overall better at classifying radiomic features than all other models within the training dataset (Figure 3a). The only exceptions were in gradient boosted trees that had better performance in four radiomic features (differences below 0.026 AUC) and random forest in one radiomic feature (0.012 AUC difference). In testing, neural networks had 0.42–0.89 AUC, 0.45–0.94 accuracy, and 0.09–0.98 average precision across all radiomic feature classifications (Supp. Fig. S4). A subset of 13 radiomic features had at least 0.70 testing AUC and was subsequently selected for interpretation. Figure 3b depicts the neural network’s generalizability to classify the aforementioned 13 radiomic features in the testing dataset.

**Figure 3:**
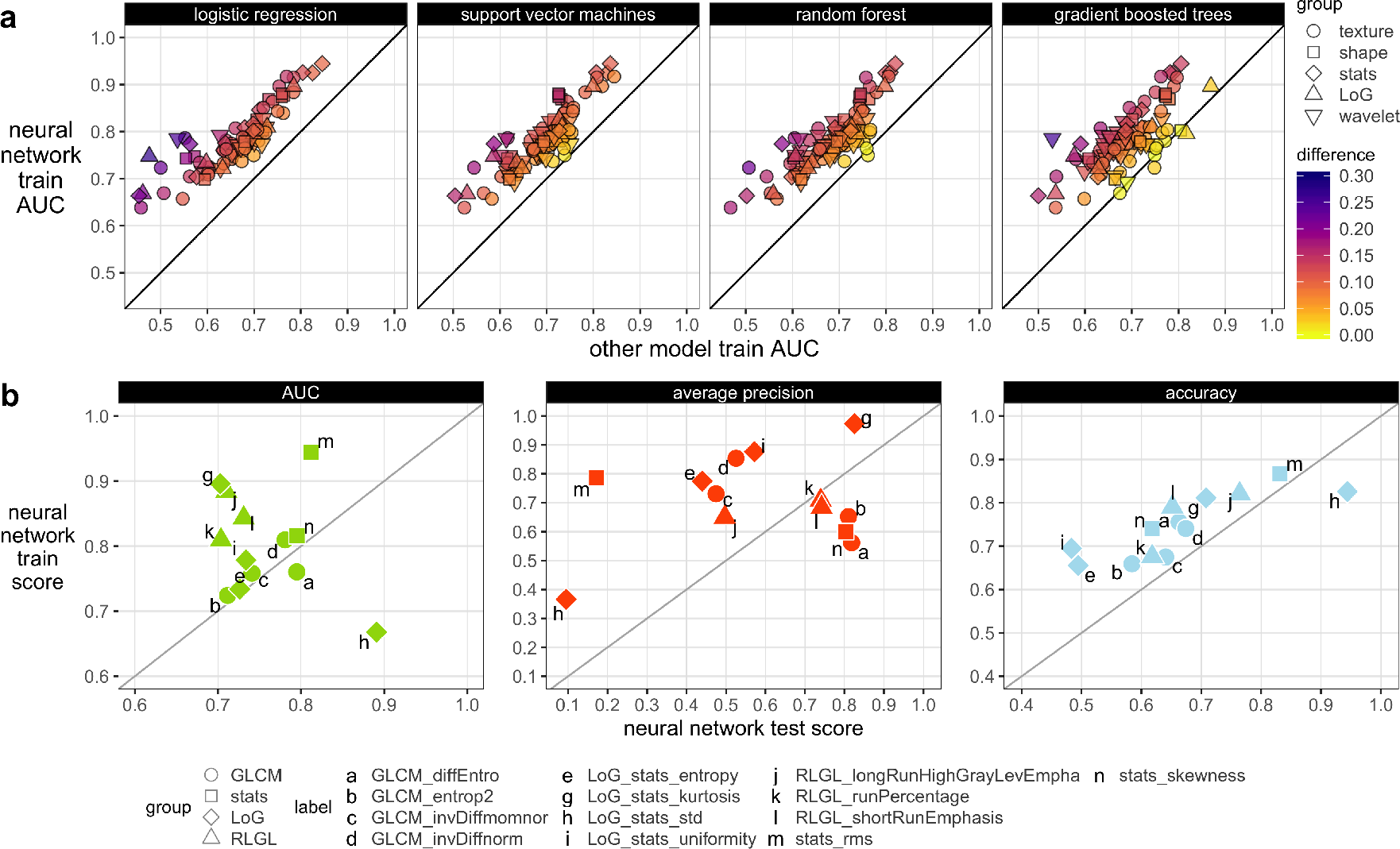
Radiogenomic modeling performance (**a**) between neural networks and other models in training, and (**b**) neural network performance in training and testing datasets of the 13 radiomic features selected for further analysis.

#### 3.2.2 Interpretation of radiogenomic associations and comparison with prior modules

Figure 4 illustrates the top GO gene sets associated with predicting each radiomic feature. Overall, gene masking suggested each radiomic feature is associated with a unique gene expression profile driven by different biological processes as none of the radiomic features had similar scores across all gene sets. Some gene sets were better for predicting one radiomic feature but not another. For example, the top two gene sets for predicting `RLGL_longRunHighGrayLevEmpha`, an imaging texture, were `regulation of syncytium formation by plasma membrane fusion` and `pyrimidine nucleotide salvage` had test AUCs above 0.75 but for all other 12 radiomic features these two gene sets were below 0.70 AUC and 0.65 AUC, respectively. Conversely, there were gene sets that could predict multiple radiomic features at once. For example, response to tumor necrosis factor was predictive of `GLCM_entrop2` (0.78 AUC, 0.81 AP) and `RLGL_runPercentage` (0.76 AUC, 0.76 AP) in testing. Hallmark gene sets were also applied to the radiomic models in gene masking analysis (Supp. Fig. S6) but were not as predictive as the GO gene sets.

**Figure 4:**
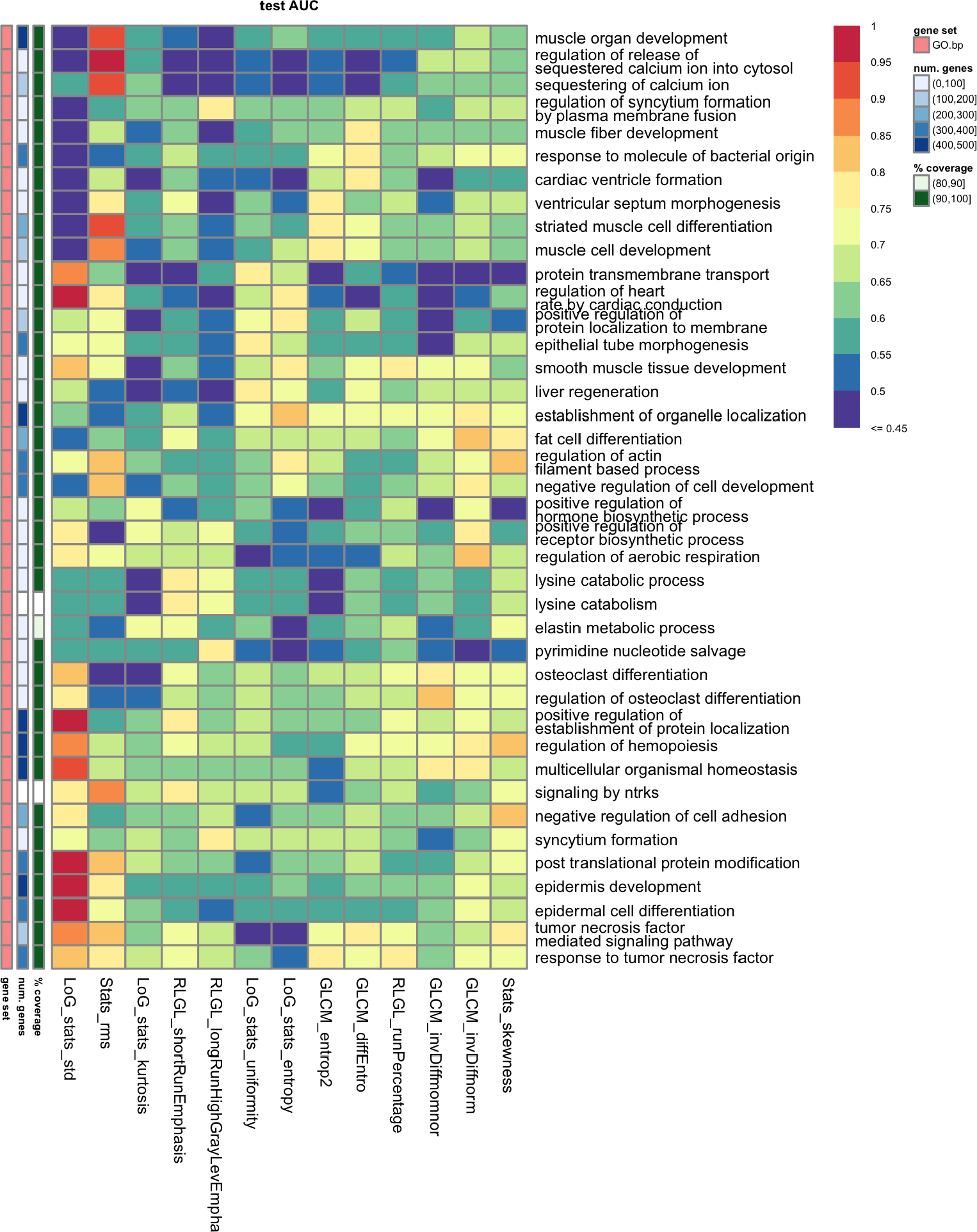
Gene masking of the radiogenomics models with biological processes from Gene ontology (GO). The top three ranked by test AUC for each radiomic feature are shown. There are 40 gene sets.

Radiogenomic associations were summarized for radiomic features related to histogram statistics of either the tumor mask, the transformation of the mask (LoG features), and textures of tumors in Table 2. The radiomic feature, `LoG_stats_std` was completely predictable in the test cohort with the gene set `post-translational protein modification` and the gene sets related to the epidermis development. Processes involving immune system and cardiac system were top predictors for several radiomic features. Many gene sets were related to cell development, varying from muscle, liver, epidermis, fat cell, and renal gene sets. AKT signaling, a targeted pathway in NSCLC therapy [19] was somewhat predictive in `RLGL_longRunHighGrayLevEmpha` with 0.76 AUC but a lower 0.54 average precision in testing. TNF was associated with `RLGL_runPercentage` and `GLCM_entrop2`. Rho signaling, associated with tumor suppressor activity and another targeted pathway in NSCLC [20, 21] was associated with `GLCM_diffEntro` (0.75 AUC; 0.78 AP) and `GLCM_invDiffmomnor` (0.79 AUC; 0.55 AP). While `GLCM_invDiffmomnor` and `invDiffnorm` were correlated and clustered together (Supp. Fig. S1), the two radiomic features had differing gene sets.

**Table 2:**
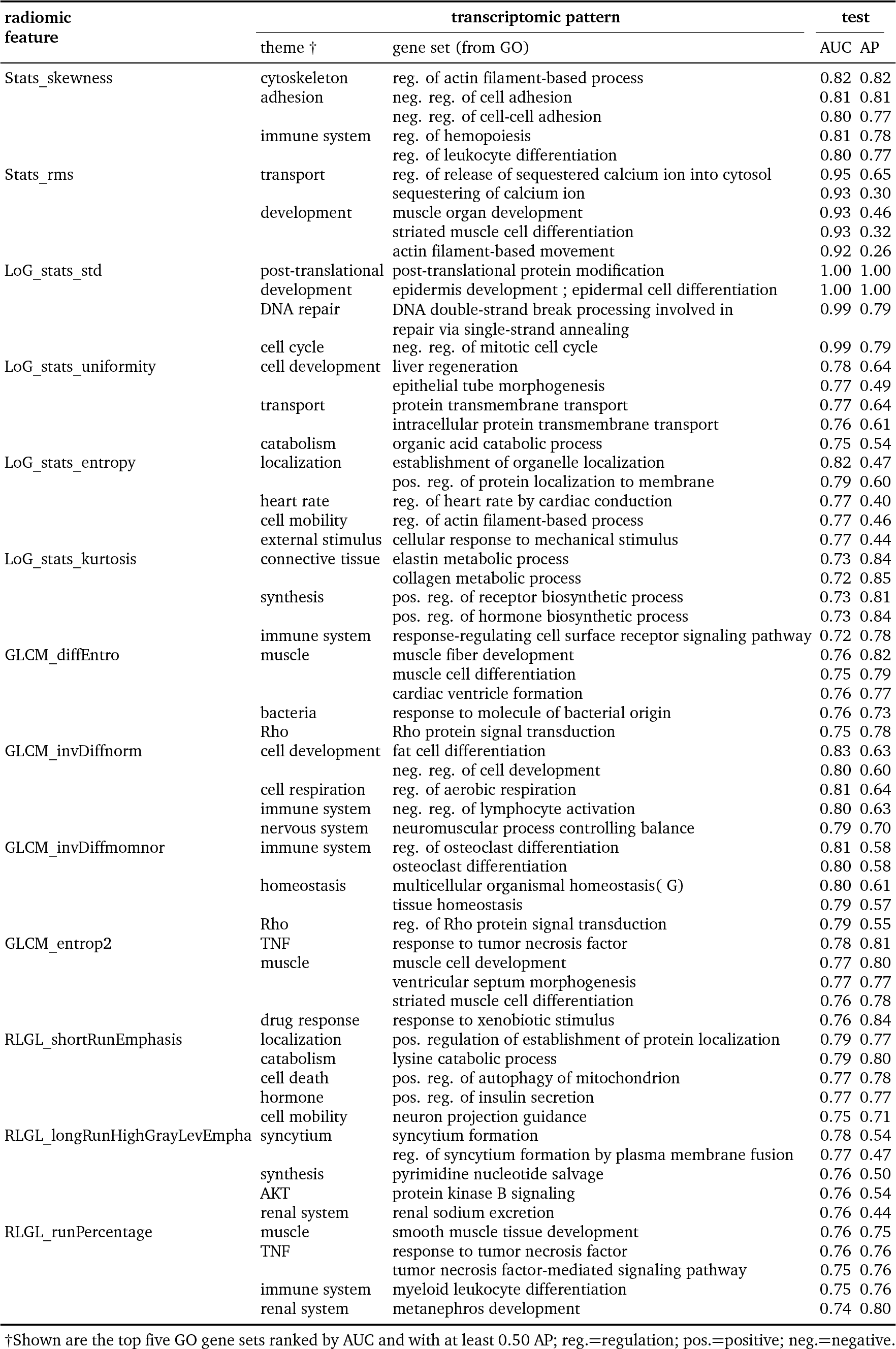
Summary of predictive gene expression patterns in NSCLC radiomic features.

Radiogenomic modules, defined as a set of correlated radiomic features and gene expressions were previously found in the same dataset [11]. Notably, not all the radiomic features were in those reported in their study’s radiogenomic modules. Here, the radiomic models were masked with the same Reactome gene sets from MSigDB v4.0 (Supp. Fig. S7). Supp. Table S3 summarizes the overlapping radiogenomic associations found in this study compared to [11]. The best agreement was between `LoG_stats_entropy` and module 13, where three of the pathways in the module were also among the top ten most predictive gene sets in our radiogenomic model. Other comparisons did not have overlapping associations. For example, the authors reported a radiogenomic association between `GLCM_diffEntro` and the five pathways in module 2, while we found the most predictive pathway in module 2 was ranked 197 out of the 664 Reactome pathways used in gene masking. Thus, our model suggested many other pathways were more predictive of the radiomic feature than the five pathways in module 2.

## 4 Discussion

We demonstrate the ability of deep neural networks to learn associations between radiomic features or clinical traits and gene expressions using two NSCLC cohorts. The datasets had a relatively large training dataset of 262 patients, and another dataset of 89 patients that allowed us to validate the generalizability of our neural network models. We show neural networks outperformed other machine learning models and had test performances above 0.70 AUC in predicting thirteen radiomic features and histology. We interpreted the models using gene masking and identified specific sets of gene expressions that were indicative of a trait or feature. Together, these results suggest potential biological associations exist to explain the differences among histology classes and CT imaging characteristics of NSCLC patients.

Previous radiomic and radiogenomic efforts have recently been studied in NSCLC [12, 22, 23, 24, 11, 25, 9]. Molecular statuses are predicted using radiomic or other imaging features in [12, 22, 24, 25]. A recent work used the same dataset of 89 to train models to predict immune cell gene signatures of NSCLC tumors using CT radiomic features [26]. Compared to these works, our model uses whole gene expression profiles to map to radiomic features. While deep neural networks are used to map CT image patches to tumor gene expressions in [9], the study does not report specific radiogenomic associations. Most related is the work from [11] (the source of the two NSCLC datasets), who use Gene Set Enrichment Analysis and the Iterative Signature Algorithm, a correlation and biclustering method to group radiomic features with Reactome gene sets (i.e., pathways) into radiogenomic modules. The top ten most pre-dictive Reactome pathways for a radiomic feature in our models have some overlap with the radiogenomic modules created in [11] but this is often not the case. These results indicate the studies identified different radiogenomic associations in the same data. Differences in radio-genomic associations between our work and these authors [11] may relate to differences in methodologies. Most importantly, our study assessed entire gene profiles, whereas their study assessed correlations between sets of radiomics features and sets of genes. As well, we did not predefine how may associations may be found prior to mapping or modeling; their study attempted to assess 20 radiogenomic modules. Finally, we used a classification (prediction) approach, whereas they used a correlation matrix to identify associations. Additionally, radiomic signatures are reportedly predictive of survival in [23, 11]. It is a part of future work to evaluate if radiogenomic relationships found in our neural network models can predict patient survival.

Compared to our prior radiogenomic work in glioblastoma (a high-grade brain tumor), NSCLC is a cancer with multiple stages and histology types. We further explore the capability of neural networks to map transcriptomes to other relevant patient imaging features by training models to predict stage and histology. While neural networks are better at predicting stage and histology in the training dataset compared to other classifiers, stage is poorly estimated in the testing dataset. However, the histology neural network has 0.77 test AUC when averaged across each histology type.

Several prior works for predicting NSCLC histology using radiomics have been based on differentiating adenocarcinoma from squamous cell carcinoma. For example, using five radiomic features a study trained a Naive Bayes model to distinguish adenocarcinoma from squamous cell carcinomas and achieved a 0.72 AUC in a test set of 152 patients [27]. In another similar study, a radiomic signature was able to distinguish adenocarcinoma from squamous carcinomas using a logistic regression model and 129 patients; the authors reported a 0.893 AUC in a test subset of 48 patients [28]. More recent work reported logistic models that achieved 0.694, 0.780, 0.800, and 0.923 AUC for clinical, standard CT features, radiomics, or all three to respectively predict adenocarcinoma versus squamous cell carcinomas [29]. The AUC scores were observed in a cohort of 93 patients but are likely overly optimistic given there was no validation or test set to evaluate their models. In contrast, our neural network model uses gene expression profiles to predict histology. In a test set of 89 patients, our model achieves a 0.86 test AUC when estimating adenocarcinoma versus all other (squamous and otherwise) histology types and a 0.91 test AUC when estimating squamous versus all other types (adenocarcinoma and otherwise).

Additionally, there is a notable difference in our models’ performances when predicting histology compared to stage. The difference is likely because staging is based on factors such as tumor size, location, and spread to lymph nodes or metastasis in other sites, which is information that may not be readily seen in gene expressions of the collected tumor tissue. On the other hand, histology classification is based on the molecular and physical characterization of a collected sample, which is likely more related to the transcriptome profiling of the tumor. Subsequently, we extract the learned associations between gene expression and histology types with gene masking and compare against two previous studies who report gene expression signatures to predict NSCLC histology [16, 17]. The studies’ gene signatures are predictive in our models as well and indicate neural networks have found similar associations in prior work. Other gene sets such as hypoxia (200 genes) and angiogenesis (36 genes) Hallmark genes sets and the neuronal (156 genes) GO gene set are associated with histology in our model and may have the potential to automate/standardize the histology determination process.

This study has several notable limitations. The retrospective datasets used in this analysis were from two different sources with varying imaging protocols, which can add variance to computationally extracted imaging features such as radiomics. A majority of patients were early stage cancers and thus late stage patients were left out of the radiogenomic associations. While sample size is an inherent concern of radiogenomic analysis, the dataset is one of the few publicly available. Still, radiomic features are difficult to interpret, and researchers are currently attempting to understand their correlation with tumor biology and other clinical traits. Radiomic features were binarized using clustering, and radiomic features with a minority class below 10% were removed. The tumor tissue samples used for transcriptome profiling are limited in that only one sample was acquired per patient and may not fully capture tumor heterogeneity. These factors make it challenging to validate the relationships in the radiogenomic neural networks. The reported findings are hypotheses about radiogenomic associations, are not proven here, and would require cell or animal studies.

In future work, the issues may be addressed by the standardization of imaging protocols to allow consistent comparison of imaging features and maps such as genomic atlases to better characterize whole tumors. A larger sample size could result in a more complete representation of the general population and allow modeling of radiomic features as continuous outputs. The ranked gene sets are interpreted based on GO descriptions. A more quantitative approach to compare the ranked gene sets between radiomic features, such as semantic similarity of GO terms [30], and a sensitivity analysis of resultant radiogenomic associations are needed. Outside of the transcriptome, there are likely other contributing factors to tumor imaging features, such as other molecular data (e.g., gene mutations and methylation) and patient covariates (e.g., smoking status). These factors could either be incorporated into the modeling process or used to stratify analyses. Additionally, the impact of knowing such radiogenomic associations at the time of tumor biopsy in relation to survival or treatment projection would be highly beneficial.

## 5 Conclusion

In this study, we present a machine learning approach based on neural networks for mapping gene expressions to radiomic features or clinical traits for patients with non-small cell lung cancer. Our models are evaluated through previously published training and testing datasets to show such neural networks are capable of modeling high-dimensional gene expression to pre-dict tumor imaging features and lung cancer histology. We further interpret the models through gene masking and report the learned relationships between gene expressions and a radiomic feature or gene expressions and a histology type, which include both new and previously reported findings in prior works. The predictive relationships found in this work could be further studied to develop automated classification of histology, non-invasive imaging surrogates for gene expressions, or individualized patient prognosis or treatment therapies.

## 6 Supporting Information

## Supplemental materials

### 1 Supplemental methods

A total of 101 radomic features were considered in this study out of the 636 from [11]. These included: 10 shape, 22 GLCM, 7 GLSZM, 10 RLGL, 12 stats, 9 LoG, and 31 wavelet features that had at least 10% of the training dataset with the least frequent class. LoG and wavelet features were selected from “LoG_sigma_0_5_mm_2D” and the “HHH” wavelet decomposition; the rest could be explored in future studies. Radiomic features not defined in [11, 12] were not considered for model interpretation and survival analysis.

Parameters for class saliency values were set to default in `keras-vis 0.4.1` [31], except for the input range (determined by the gene expression range in the dataset) and `backprop_modifier` set to guided.

### 2 Supplemental tables and figures

#### 2.1 Dataset

**Table S1:** Patient characteristics, estimated or replicated from [11].

|  |  | training<br>n=262 | testing<br>(n=89) |
| --- | --- | --- | --- |
| source | country | USA | Netherlands |
| gender | male | 100 (38%) | 59 (66%) |
|  | female | 124 (47%) | 28 (32%) |
|  | n/a | 38 (15%) | 2 (2%) |
| histology | adenocarcinoma | 129 (49%) | 42 (48%) |
|  | squamous | 61 (23%) | 33 (38%) |
|  | other | 34 (13%) | 12 (14%) |
|  | n/a | 38 (15%) | 0 (0%) |
| stage | I | 123 (47%) | 39 (44%) |
|  | II | 35 (13%) | 26 (29%) |
|  | III | 46 (18%) | 12 (14%) |
|  | other | 20 (8%) | 10 (11%) |
|  | n/a | 38 (14%) | 2 (2%) |
| smoking status | current | 66 (25%) | n/a |
|  | former | 141 (54%) | n/a |
|  | none | 17 (7%) | n/a |
|  | n/a | 38 (14%) | n/a |
| tumor site | primary | 224 (86%) | 87 (98%) |
|  | n/a | 38 (14%) | 2 (2%) |
| status | alive | 134 (51%) | 41 (46%) |
|  | deceased | 90 (35%) | 46 (52%) |
|  | n/a | 38 (14%) | 2 (2%) |
| follow-up | median months | 32 | 31 |

**Figure S1:**
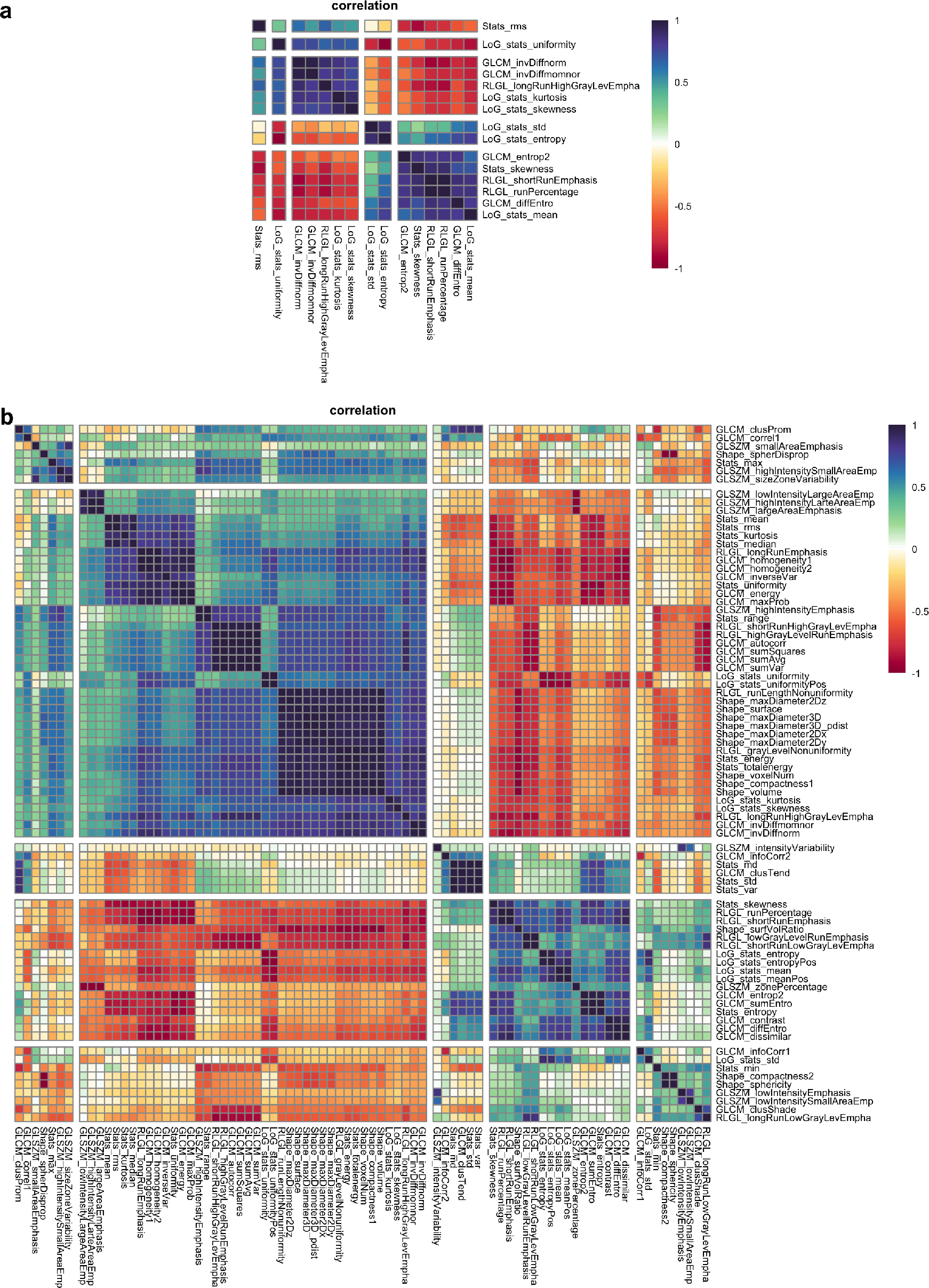
Correlation between radiomic features in Dataset1, the training dataset. (a) The subset of radiomic features selected based on classification performance in testing. (b) All radiomic features used in this study (n=101). Features were clustered with `pheatmap in R` using the Spearman rank correlation and default parameters.

**Figure S2:**
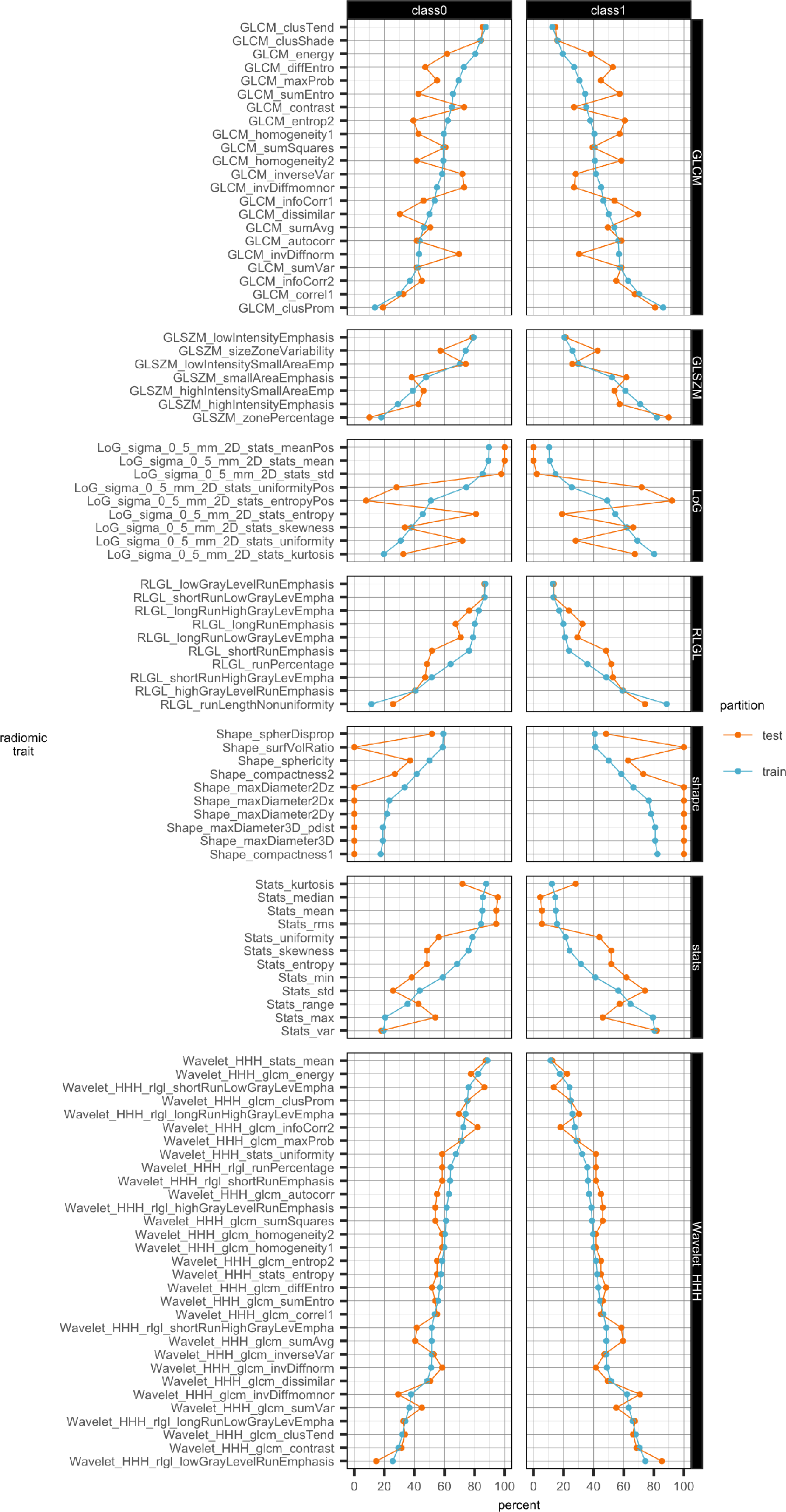
Class distributions of radiomic features after clustering into two classes.

#### 2.2 Modeling

**Figure S3:**
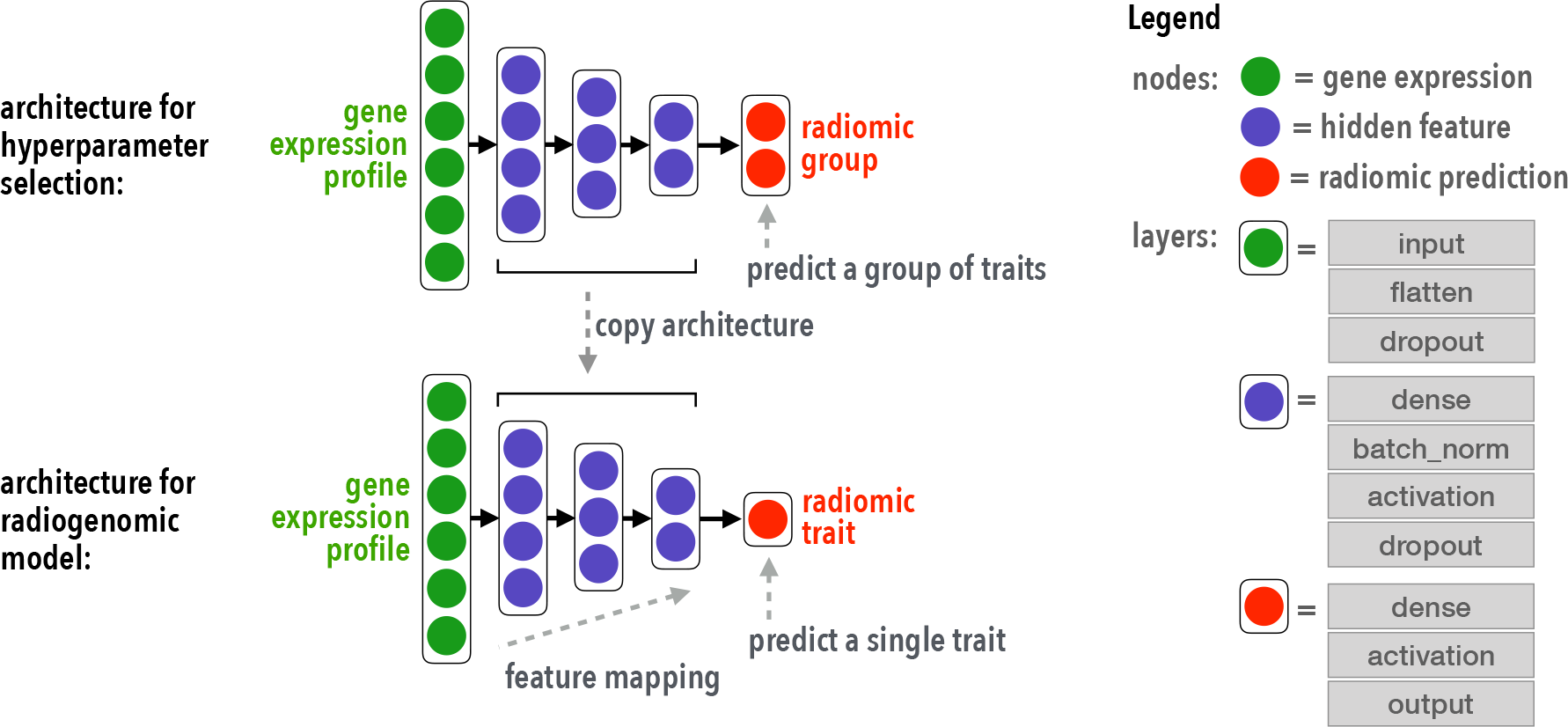
The architecture and hyperparameter tuning of a radiogenomic neural network.

**Table S2:**
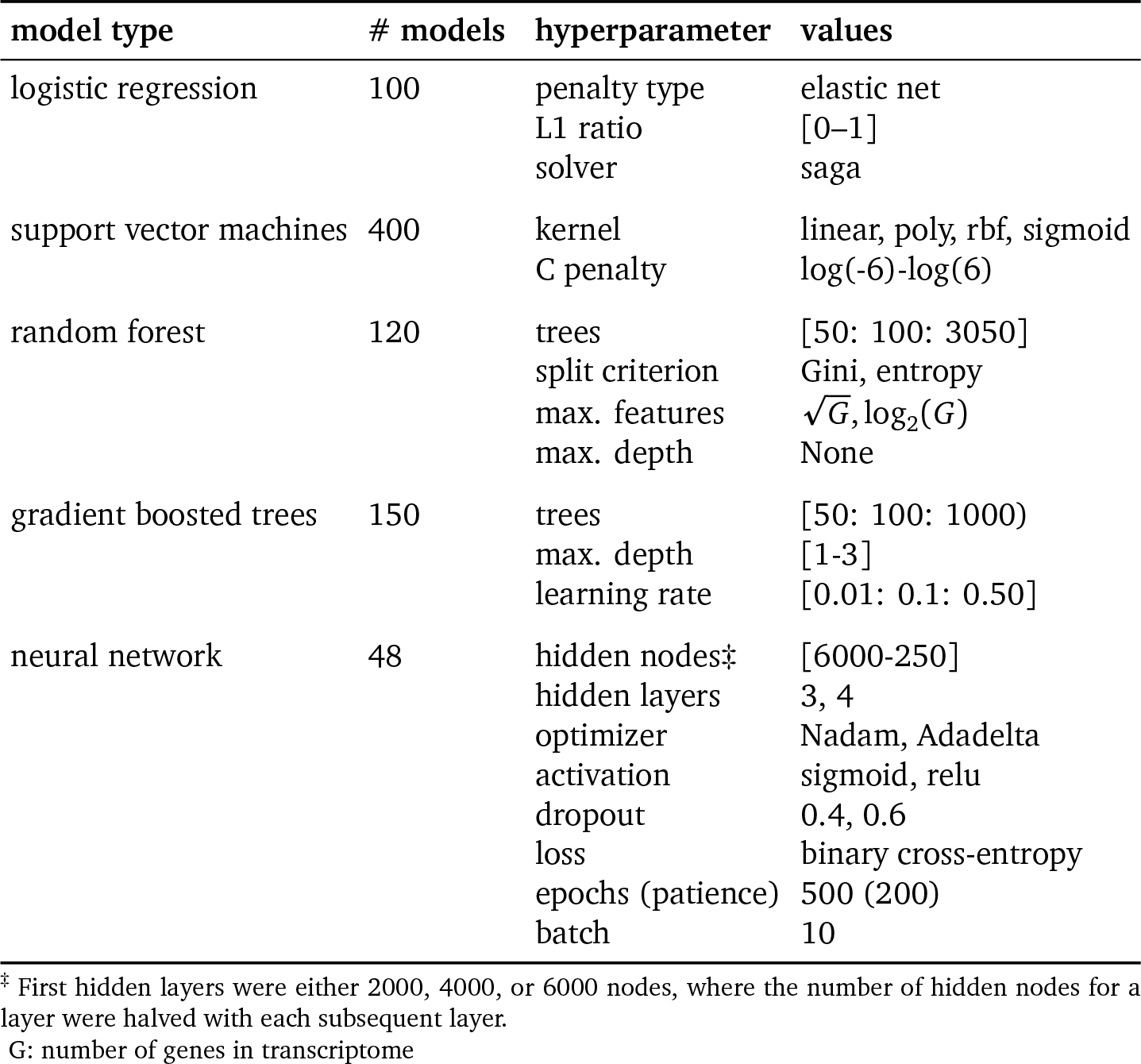
NSCLC radiogenomic models and hyperparameters. Grid search was used for selecting hyperparameters

| model type | # models | hyperparameter | values |
| --- | --- | --- | --- |
| logistic regression | 100 | penalty type<br>L1 ratio<br>solver | elastic net<br>[0–1]<br>saga |
| support vector machines | 400 | kernel<br>C penalty | linear, poly, rbf, sigmoid<br>log(-6)-log(6) |
| random forest | 120 | trees<br>split criterion<br>max. features<br>max. depth | [50: 100: 3050]<br>Gini, entropy<br>$\sqrt{G}$, $\log_2(G)$<br>None |
| gradient boosted trees | 150 | trees<br>max. depth<br>learning rate | [50: 100: 1000)<br>[1-3]<br>[0.01: 0.1: 0.50] |
| neural network | 48 | hidden nodes‡<br>hidden layers<br>optimizer<br>activation<br>dropout<br>loss<br>epochs (patience)<br>batch | [6000-250]<br>3, 4<br>Nadam, Adadelta<br>sigmoid, relu<br>0.4, 0.6<br>binary cross-entropy<br>500 (200)<br>10 |

‡ First hidden layers were either 2000, 4000, or 6000 nodes, where the number of hidden nodes for a layer were halved with each subsequent layer.
G: number of genes in transcriptome

**Figure S4:**
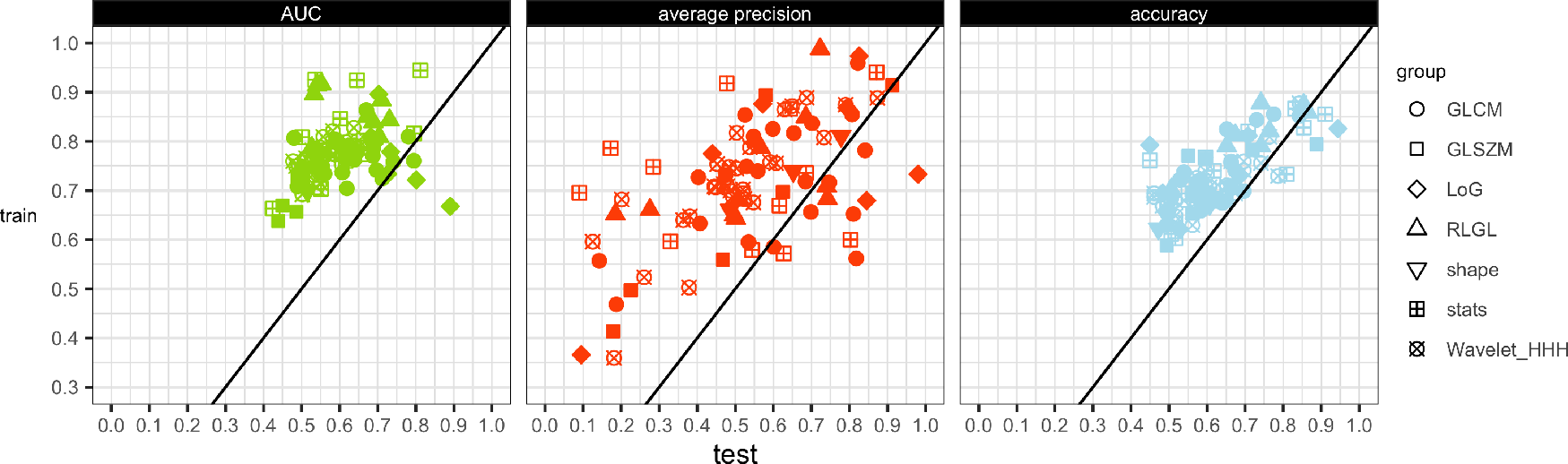
Classification performance of radiogenomic neural networks used to predict 101 radiomic features in training and testing. Not all radiomic features had both binary labels in testing; therefore some metrics are missing. Accuracy was defined at a 0.5 threshold of predictions.

#### 2.3 Gene masking

**Table S3:** A comparison of the learned radiogenomic associations extracted from our neural networks and the modules identified in [11]. Each module consisted of a set of Reactome pathways and a set of imaging features. Shown are the modules that included the radiomic features used in this study. If any module’s set of pathways was ranked among the top 100 in gene masking, the top three pathways were listed.

| radiomic trait | Reactome pathway | this study |  | [11] |  |
| --- | --- | --- | --- | --- | --- |
|  |  | test AUC | rank† | module | # pathways |
| GLCM_diffEntro | cross presentation of soluble exogenous antigens endosomes | 0.58 | 197 | 2 | 5 |
|  | phase II conjugation | 0.69 | 9 | 12 | 35 |
|  | regulation of mitotic cell cycle | 0.66 | 21 |  |  |
|  | ABCA transporters in lipid homeostasis | 0.65 | 28 |  |  |
| LoG_stats_std | regulation of ornithine decarboxylase (ODC) | 0.94 | 15 | 2 | 5 |
|  | cross presentation of soluble exogenous antigens endosomes | 0.82 | 89 |  |  |
|  | antigen processing cross presentation | 0.80 | 106 |  |  |
|  | cholesterol biosynthesis | 0.75 | 150 | 6 | 7 |
|  | signaling by TGF beta receptor complex | 0.83 | 81 | 8 | 17 |
|  | elongation arrest and recovery | 0.82 | 92 |  |  |
|  | mRNA splicing | 0.75 | 142 |  |  |
| LoG_stats_entropy | elongation arrest and recovery | 0.73 | 1 | 7 | 8 |
|  | mitochondrial protein import | 0.63 | 74 |  |  |
|  | RNA pol III chain elongation | 0.58 | 180 |  |  |
|  | elongation arrest and recovery | 0.73 | 1 | 13 | 26 |
|  | RNA pol II pre transcription events | 0.69 | 7 |  |  |
|  | formation of RNA pol II elongation complex | 0.69 | 9 |  |  |
| Stats_skewness | antigen processing cross presentation | 0.62 | 139 | 2 | 5 |

† All Reactome pathways were ranked by test AUC.

**Figure S5:**
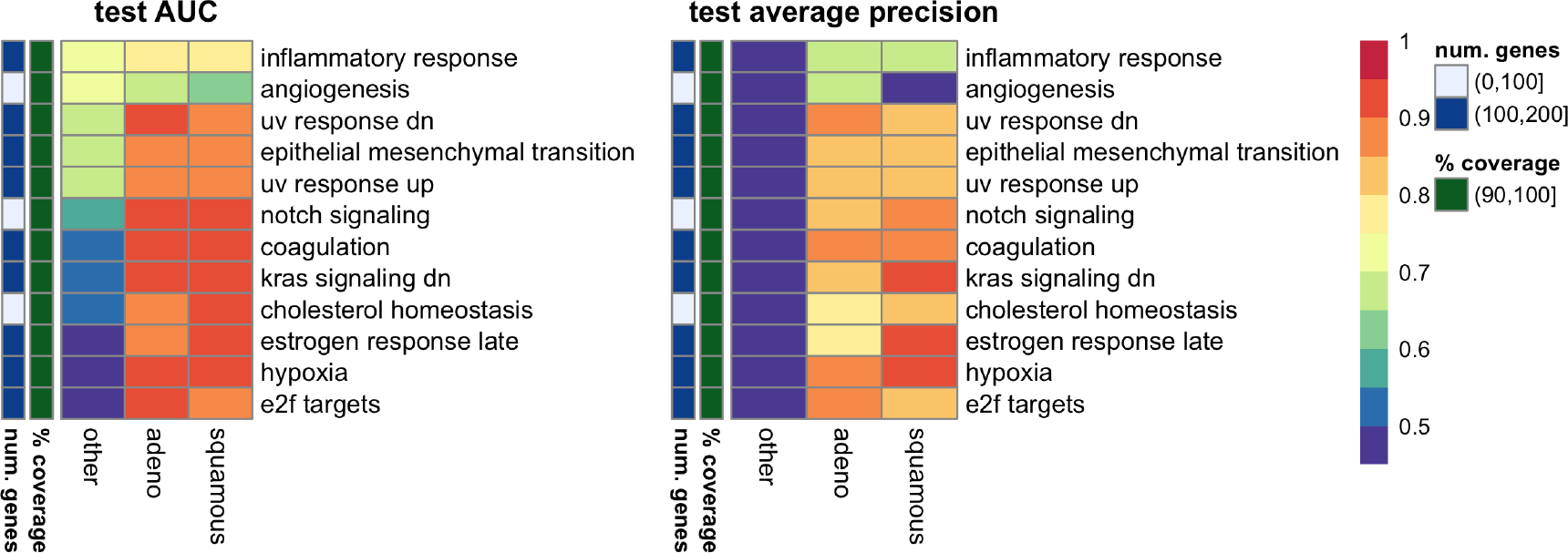
Hallmark gene set masking in the histology model. The top five ranked by AUC for each histology type were shown. There are twelve hallmarks.

**Figure S6:**
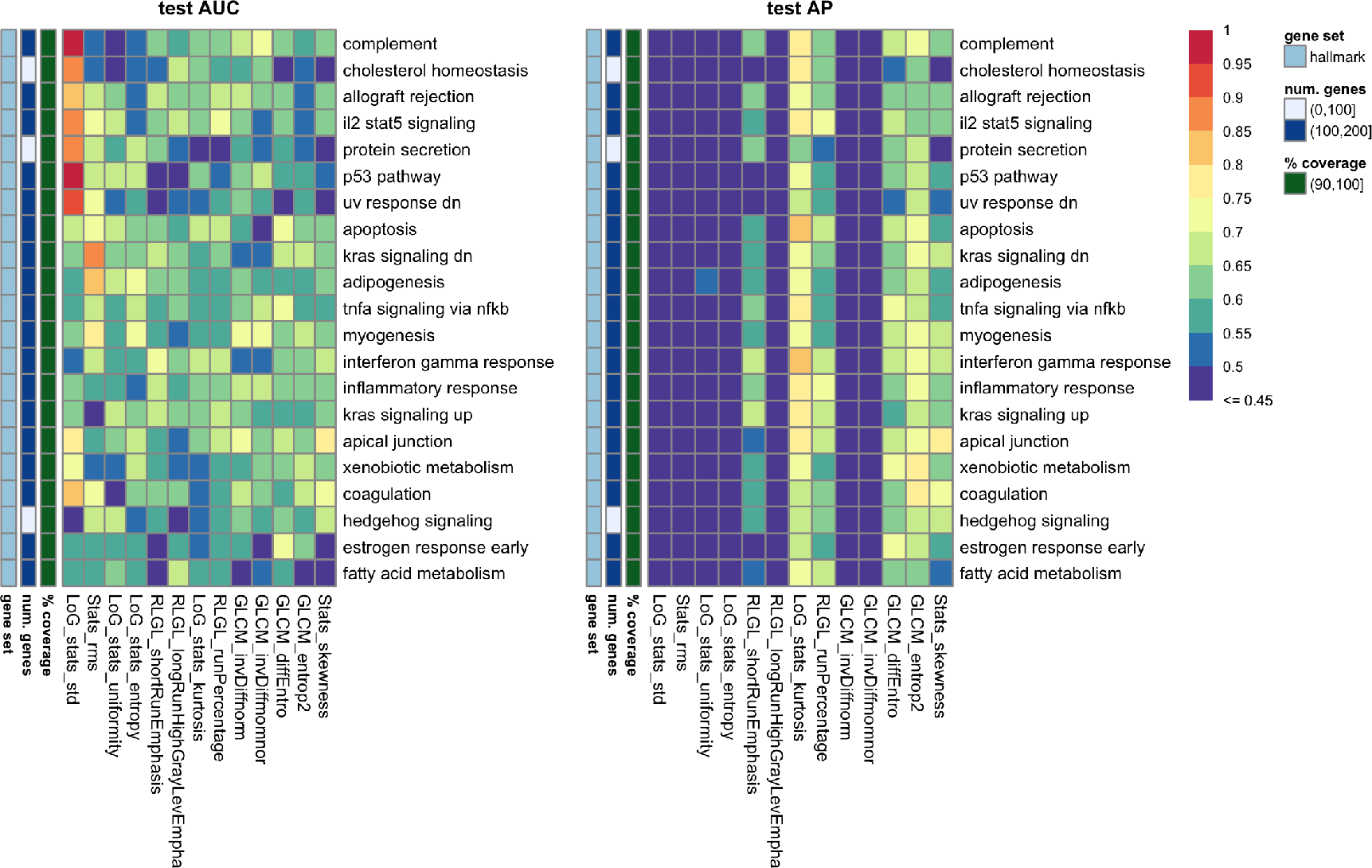
Hallmark gene set masking in the histology model. The top three ranked by AUC for each radiomic trait were shown. There are 21 hallmarks.

**Figure S7:**
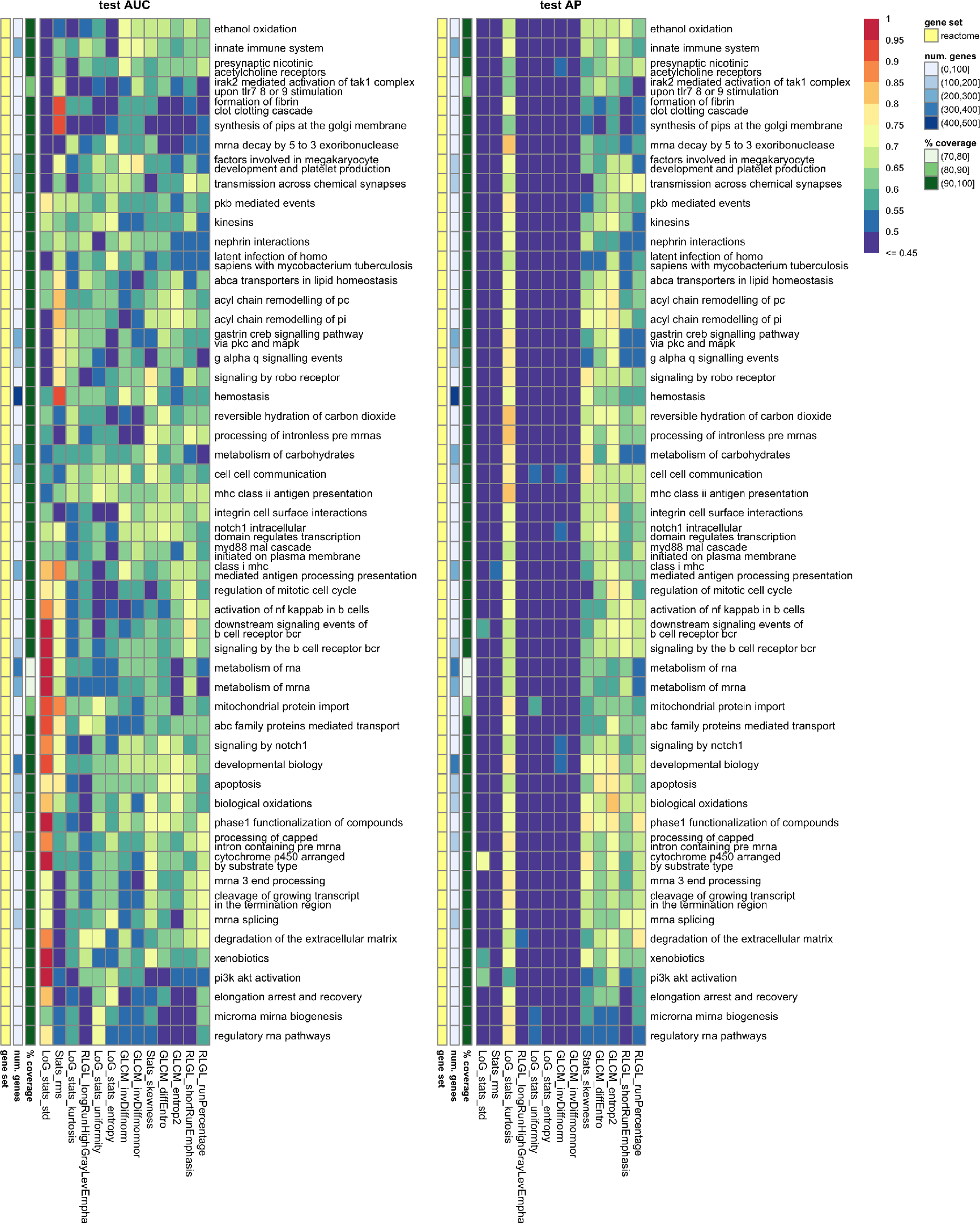
Reactome gene set masking in the histology model. The gene sets are from MSigDB v4.0 to match the pathways previously used in [11] to allow a comparison. The top five ranked by AUC for each histology type were shown. There are 53 pathways shown. There were 674 pathways total and 669 pathways with less than 500 genes.

## Notes

**Conflicts:** The authors declare no potential conflicts of interest.

